# Contrasting adaptation and optimization of stomatal traits across communities at continental-scale

**DOI:** 10.1101/2021.11.30.470674

**Authors:** Congcong Liu, Lawren Sack, Ying Li, Nianpeng He

**Author notes:** Correspondence author Nianpeng He.

## Abstract

The maximum stomatal conductance (*g*), a major anatomical constraint on plant productivity, is a function of the stomatal area fraction (*f*) and stomatal space-use efficiency (*e*). However, *f* and *g* have been considered as equivalents, with *e* rarely considered, and their adaptation to the environment and their regulation of ecosystem productivity are unclear. Here, we analyzed the community-weighted mean, variance, skewness, and kurtosis of stomatal traits from tropical to cold-temperature forests. The variance of *g* and *f* was higher for arid sites, indicating greater functional niche differentiation, whereas that for *e* was lower, indicating convergence in efficiency. Besides, when other stomatal trait distributions remained unchanged, increasing kurtosis but decreasing skewness of *g* would improve ecosystem productivity, and *f* showed the opposite patterns. These findings highlight how the relative importance and equivalence of inter-related traits can differ at community scale.

## Introduction

Stomata are micropores on the leaf surface that regulate the exchange of water vapor and CO_2_ between plants and the atmosphere (Edwards et al., 1998; HetheringtonandWoodward, 2003). Indeed, the evolution of stomata was necessary for plants to colonize terrestrial ecosystems and the diversification of stomatal traits enables plants to inhabit a wide range of environments (Haworth et al., 2011; Raven, 2002). The numbers of stomatal pores, and their area and depth determine the maximum stomatal conductance (*g*), which represent an anatomical constraint on the maximum rates of diffusion of carbon and water, and thereby their fluxes in given environments. Indeed, given that there is a close relationship between *g* and field-measured stomatal conductance (McElwain et al., 2016; Murray et al., 2019; XiongandFlexas, 2020), and *g* has been used to predict water vapor and CO_2_ fluxes (FranksandBeerling, 2009; McElwain et al., 2016; SackandBuckley, 2016). In turn, *g* can be considered a product of the fraction of leaf epidermal space that is allocated to the stomata (the stomatal area fraction; *f*) and the stomatal space-use efficiency (*e*), which is a function of stomatal size (see Methods). The *f* is more properly an index of the combined costs associated with the construction, operation and maintenance of the stomata, but it is often taken as a proxy for *g* (HollandandRichardson, 2009; Liu et al., 2018; Sack et al., 2003), especially as *g* and *f* are theoretically and empirically correlated with each other (de Boer et al., 2016; FranksandBeerling, 2009). Yet the relative importance and the equivalence of these traits have not been tested at a large scale.

The importance of *g* and its determinants is especially critical for the understanding of the adaptation of diverse species of communities across gradients of aridity. A rich literature shows contrasting trait values enables co-occurring species to exploit different resources, or the same resources on contrasting spatial or temporal scales (Gross et al., 2017; Hooper, 1998; Hooper et al., 2005), resulting in species-variation in tolerances of scarcity, e.g., drought (Grossiord, 2020), as well as facilitation (Callaway, 1995) and “selection effects”, i.e,. differential contribution to the community-weighted trait values (Loreau, 2000). Indeed, traits that contribute to resource partitioning, such as root stratification (Schwendenmann et al., 2015) or differential stomatal regulation (West et al., 2012) can contribute not only to the mechanisms by which plants tolerate drought but also can improve species-specific soil moisture status by reducing competition for water among species. As a composite stomatal trait, *g* is coordinated with other plant hydraulic traits (Sack et al., 2003). Generally, a higher *g* should benefit species under selection for high productivity or competition (SackandBuckley, 2016; Taylor et al., 2012), and thus, in communities with high water availability, we expected narrower functional niche differentiation of *g* than in communities of drier regions. Indeed, because plants can tolerate drought by maintaining low rates of water uptake and productivity as soils dry, i.e., “tolerance” and/or by achieving their growth primarily when water is available, i.e., “avoidance” (Grubb, 1998; HetheringtonandWoodward, 2003), we hypothesized that *g* values, which influence water uptake and productivity, would tend to be more variable in communities of drier regions. Notably, as *g* depends on *f* and *e*, where *g* is a proxy for the benefit of assimilated carbon, and *f* represents the cost of stomatal construction, maintenance and spatial allocation (de Boer et al., 2016), therefore *e*, which is *g*/*f*, is a benefit-cost ratio, i.e,. the maximum amount of CO_2_ that can diffuse through a unit of stomatal area per unit time. We thus hypothesized that that variability of *e* within communities would be strongly constrained under water scarcity.

Stomatal traits may be a model for plant traits that are important in determining ecosystem functions, as this role of traits has become a priority topic in ecological research (Reichstein et al., 2014). The effect of species’ traits aggregated at ecosystem scale is typically quantified using to the mass ratio hypothesis or the niche complementarity hypothesis.

According to the mass ratio hypothesis the extent to which the trait of a given species affects ecosystem properties depends on its relative contribution to the total community biomass (Garnier et al., 2004), and many studies found correlations between ecosystem functions and community-weighted mean (CWM) values of plant traits (Garnier et al., 2004; Griffin-Nolan et al., 2018; MuscarellaandUriarte, 2016). According to the niche complementarity hypothesis, resource niches may be used more completely when a community is functionally more diverse (SchumacherandRoscher, 2009), and many studies reported that ecosystem function can be predicted by niche complementarity of traits, as quantified using community-weighted variance, skewness, or kurtosis of trait values (Gross et al., 2017; Le Bagousse-Pinguet et al., 2017; Liu et al., 2020; Mensah et al., 2020; Zhang et al., 2019). Indeed, the global vegetation models predict ecosystem production based on the mean values of traits. It is still a missing picture that how trait distributions influence the prediction. Although stomatal traits are expected to influence ecosystem productivity given their essential role in controlling leaf water and CO_2_ fluxes (HetheringtonandWoodward, 2003; Wang et al., 2015), no studies have tested the relative importance of the distributions of stomatal traits (including community-weighted mean, variance, skewness, and kurtosis) in predicting ecosystem productivity across communities. We hypothesized a strong importance of these community distribution metrics for *g* and potentially for its components, *f* and *e*, for regulating ecosystem productivity at community scale.

We analyzed the community-weighted mean, variance, skewness, and kurtosis and relationships among these statistical moments, for *g, f* and *e* for 800 plant species from nine sites along a climatic gradient. We hypothesized that the community-weighted variance in *g* would increase with aridity, due to variability of *f*, rather than *e*. We also hypothesized that functional niche differentiation of *g* would be stronger for communities at higher aridity, and tested whether trait assembly of stomata followed the general assembly rule for maximization of trait diversity previously reported for drylands globally using specific leaf area and maximum plant height (Gross et al., 2017). We also hypothesized that stomatal distributions would predict differences in productivity across ecosystems.

## Results

### Relationships between stomatal trait moments and climate

Stomatal traits were closely related to temperature, precipitation, and climatic aridity (Fig. 1). Overall, the relationships of community-weighted trait means and variances with climate variables were stronger than those of community-weighted skewness and kurtosis, and the aridity index was a stronger predictor of stomatal traits than temperature and precipitation. The community-weighted means and variances of *g* and *f* were strongly positively associated with climatic aridity whereas those of *e* were negatively associated with climatic aridity (Fig. 2).

**Fig. 1.**
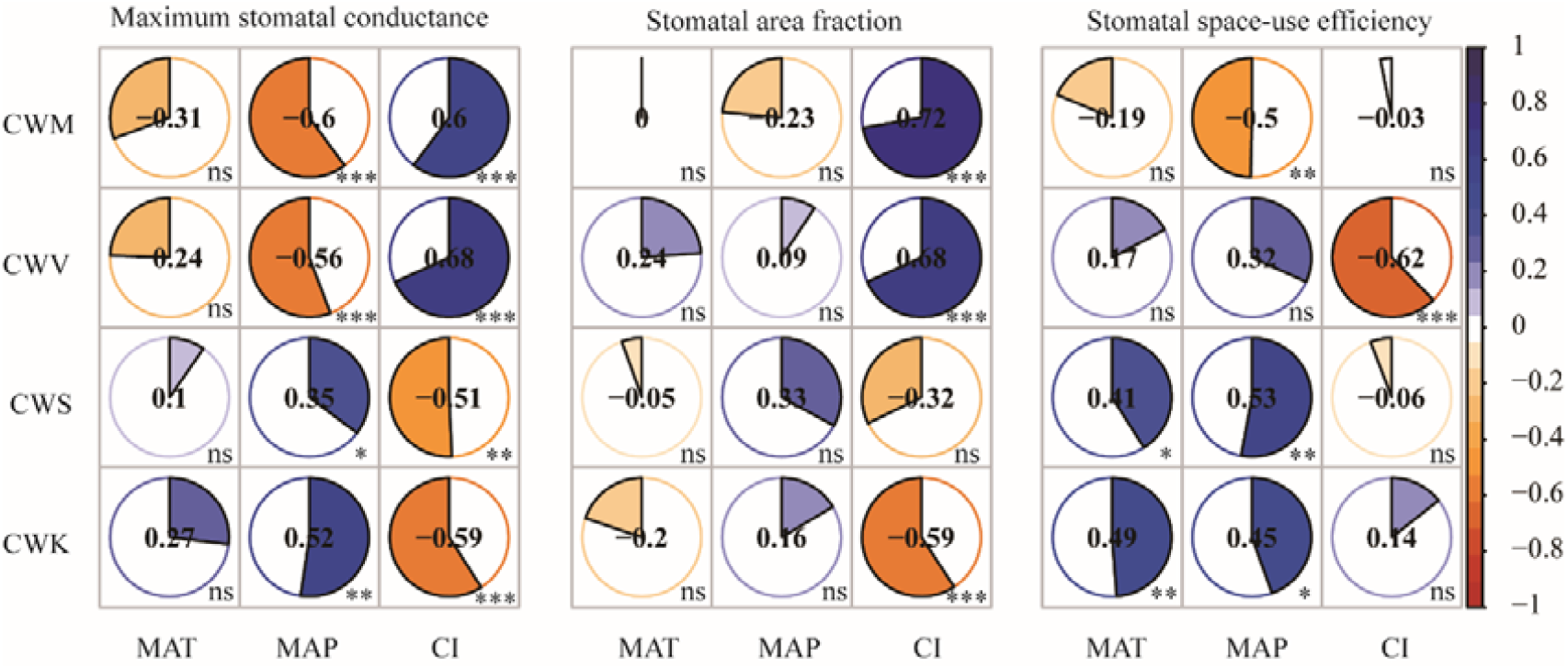
Stomatal trait moments are broadly related to climatic aridity. CWM, community-weighted mean; CWV, community-weighted variance; CWS, community-weighted skewness; CWK, community-weighted kurtosis MAT, mean annual temperature; MAP, mean annual precipitation; CI, climatic aridity index; Spearman rank correlation coefficients are shown in the panels. Fan-shaped areas are proportional to the absolute Spearman rank correlation coefficients; negative correlations are drawn with a counterclockwise fan and positive correlations with a clockwise fan. The strength of negative correlation increases from white to red, and the strength of positive correlation increases from white to blue. ns, no significance at the 0.05 level; *, *p* < 0.05; **, *p* < 0.01; ***, *p* < 0.001.

**Fig. 2.**
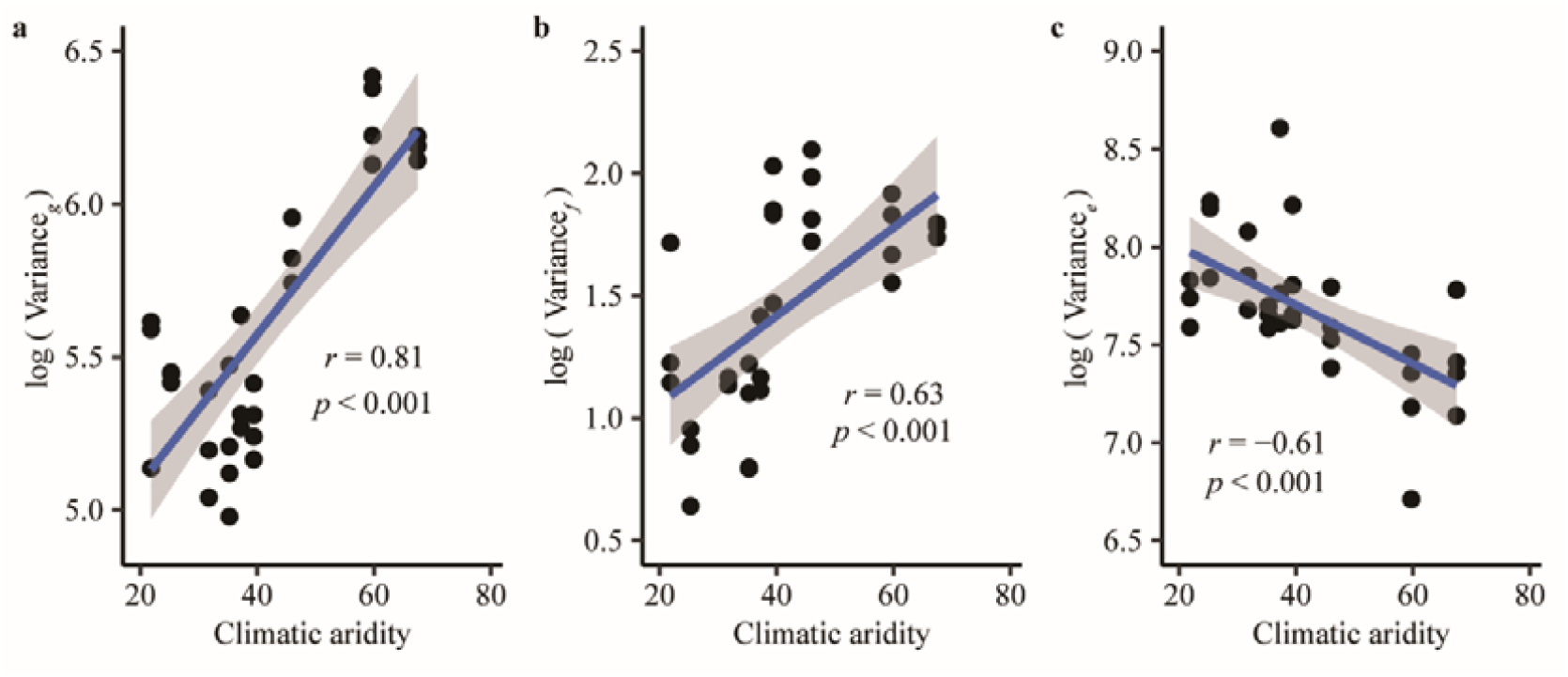
Relationships between the community-weighted variance of stomatal traits and climatic aridity. Variance, community-weighted variance. *g*, maximum stomatal conductance; *f*, stomata area fraction; *e*, stomatal space-use efficiency. The blue lines are fitted using linear regression, and shaded areas indicate the 95% confidence interval.

The correlations between community-weighted variance and kurtosis were also tested (Fig. S1). For *g* and *f*, the community-weighted variance and kurtosis were negatively correlated; such correlations were not observed with *e*. At drier sites, *g* generally showed larger variance with lower kurtosis, whereas communities of wetter sites generally had smaller variance with a wide range of kurtosis.

### Skewness-kurtosis relationships (SKR) and random expectations

In most cases, the distributions of stomatal traits differed substantially from normality (Fig. 3). The community-weighted skewness^2^ and kurtosis of these three stomatal traits were strongly positively related. The skewness and kurtosis values generated by the null model were located within the constraint triangle imposed by the inequality Kurtosis ≥ Skewness^2^ + 1. The observed empirical skewness-kurtosis relationships (SKR) for *g* deviated strongly from the predictions of the two null models, with the slopes (β) were higher and intercepts (α) lower than would be expected by chance, based on Monte Carlo analyses (Table S6). The observed kurtosis values for both *g* and *f* were significantly closer than expected by chance to the lower boundary of the mathematical constraint triangle. In other words, after controlling for the degree of skewness of *g* and *f*, observed kurtosis within communities was minimal. Skewness-kurtosis relationships for *e* did not differ statistically from those generated by the two null models; thus, the D of *e* was not smaller than expected (Fig. 3).

**Fig. 3.**
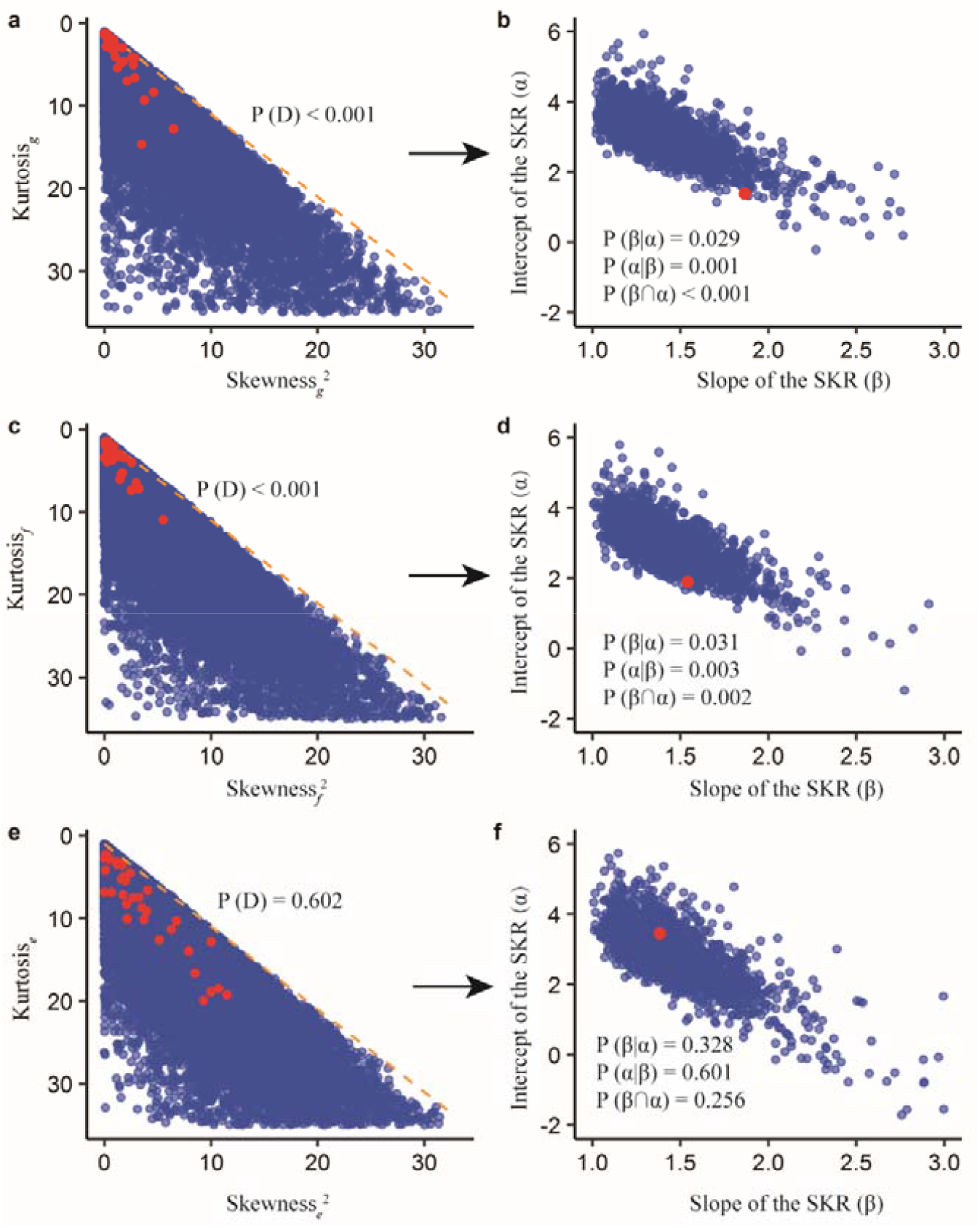
Observed skewness-kurtosis relationships (SKR) and deviation from null expectations. Skewness, community-weighted skewness; Kurtosis, community-weighted kurtosis. *g*, maximum stomatal conductance; *f*, stomatal area fraction; *e*, stomatal space-use efficiency. The red dots in the left panels represent the observed skewness and kurtosis values; blue dots in the left panels represent the skewness and kurtosis values of simulated random communities. The orange line represents *y* = *x* +1. Red/blue dots in the right panels represent the observed/random slope (α) and intercept (β) of the SKRs. We indicate the conditional pseudo P values from null model ‘richness’ for the slope β, P(β|α), the y-intercept α, P(α|β), the whole model, P(β∩α) and the distance to the lower boundary, P(D) (see Table S6 for details).

Although the skewness-kurtosis relationships of *g* and *f* cannot be explained by chance, the D of *g* and *f* was also influenced by climate. Specifically, drier communities had lower D values for *g* and *f*, while for *e* the D values showed no climatic trends (Fig. 4).

**Fig. 4.**
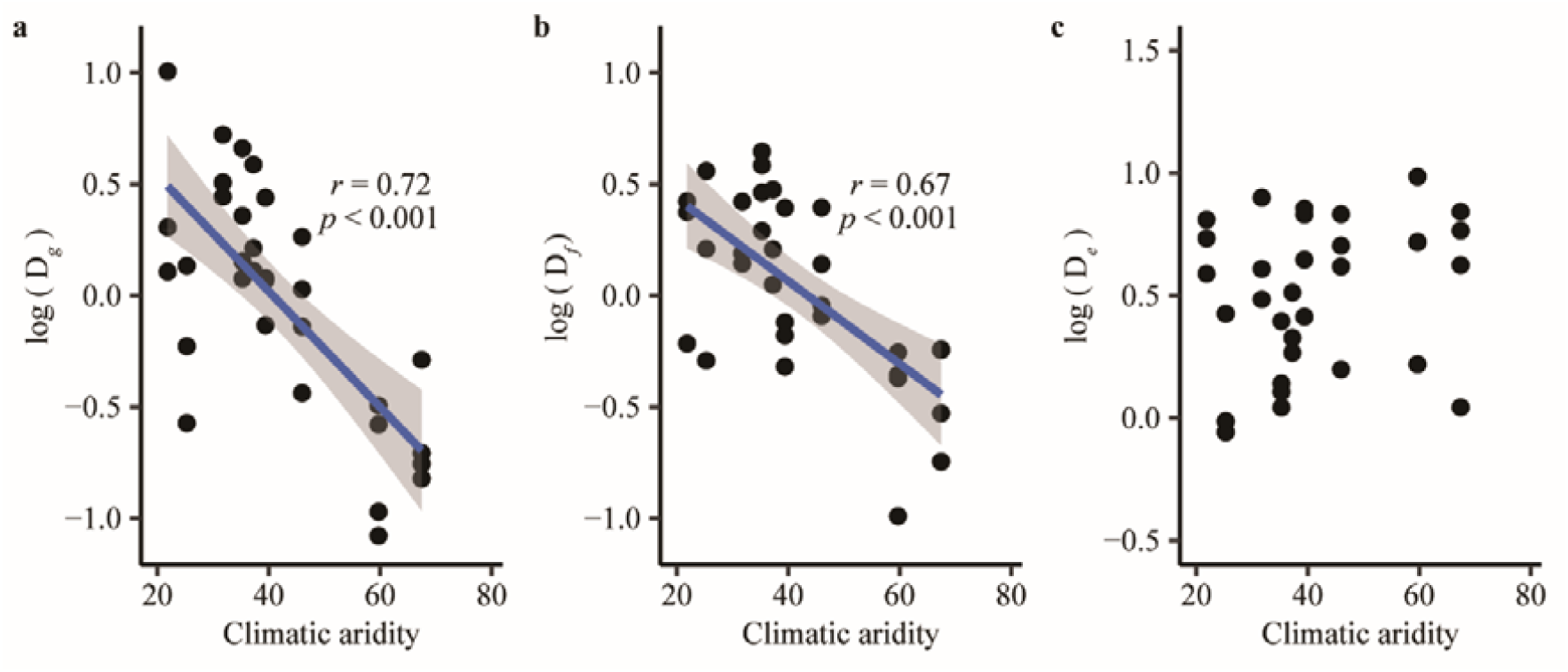
Relationships between the distance to the lower boundary (D) and climatic aridity. *g*, maximum stomatal conductance; *f*, stomatal area fraction; *e*, stomatal space-use efficiency. The blue lines were fitted using linear regression and the shaded areas indicate the 95% confidence interval.

### Stomatal trait moments and ecosystem productivity

The distributions of stomatal traits regulated ecosystem productivity (Fig. 5). The amount of variance in ecosystem productivity explained by community-weighted skewness and kurtosis was greater than that explained by community-weighted mean and variance.

**Fig. 5.**
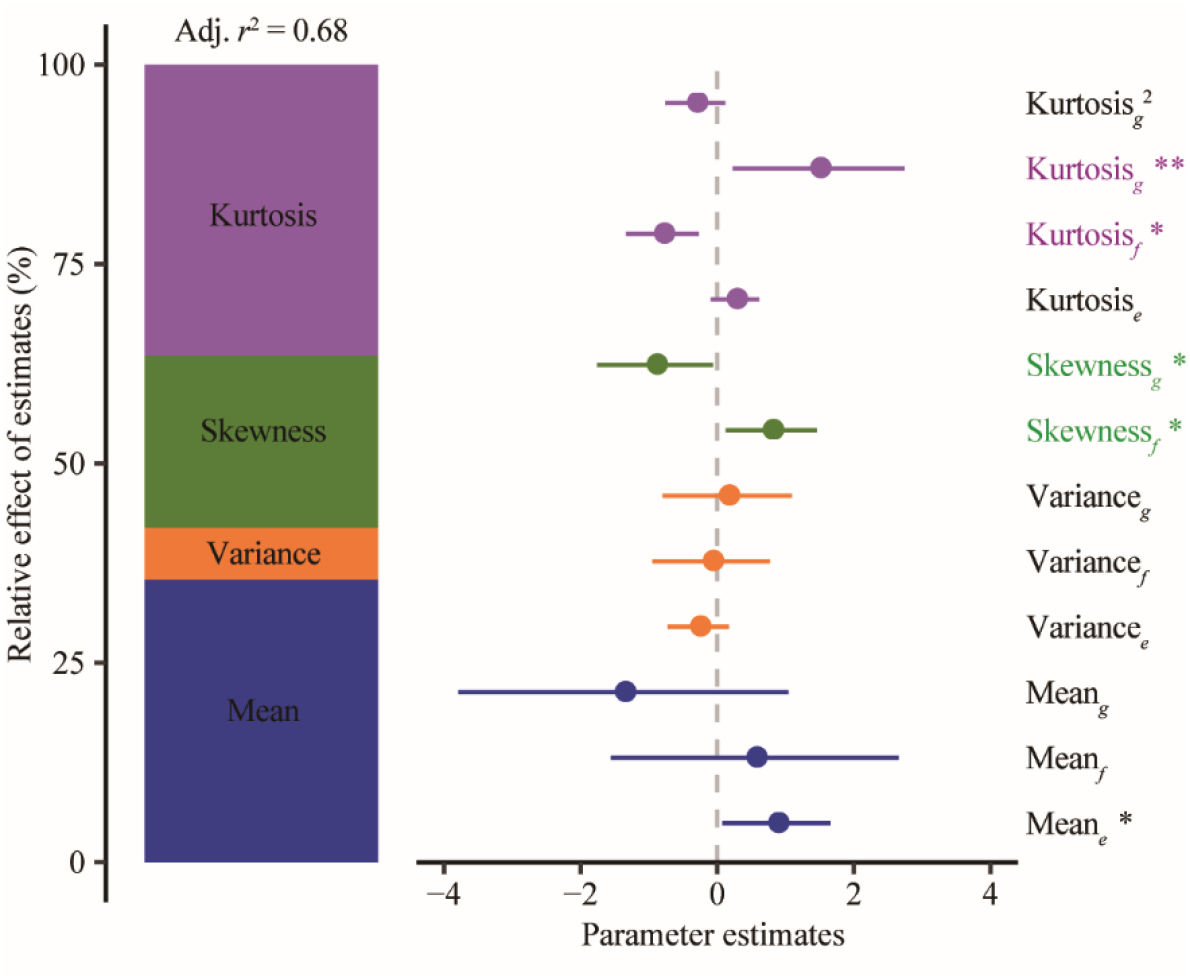
Community-weighted skewness and kurtosis of *g* and *f* showed opposite effects on ecosystem productivity. Mean, community-weighted mean; Variance, community-weighted variance; Skewness, community-weighted skewness; Kurtosis, community-weighted kurtosis. *g*, maximum stomatal conductance; *f*, stomatal area fraction; *e*, stomatal space-use efficiency. Average parameter estimates (standardized regression coefficients) of model predictors, associated 95% confidence intervals, and relative importance of each factor, expressed as the percentage of explained variance. The adjusted *r*^2^ of the averaged model and the *p* value of each predictor are given as: *, *p* < 0.05; **, *p* < 0.01. Colored labels in the right highlighted the different effects of *g* and *f* on ecosystem productivity.

Community-weighted skewness and kurtosis of *g* and *f* played different roles in optimizing ecosystem productivity: If the other independent variables were fixed, increasing the skewness of *f* but decreasing that of *g*, and increasing the kurtosis of *g* but decreasing that of *f* would improve ecosystem productivity (Fig. 6). Further, ecosystem productivity increased across communities positively with the mean of *e*.

**Fig. 6.**
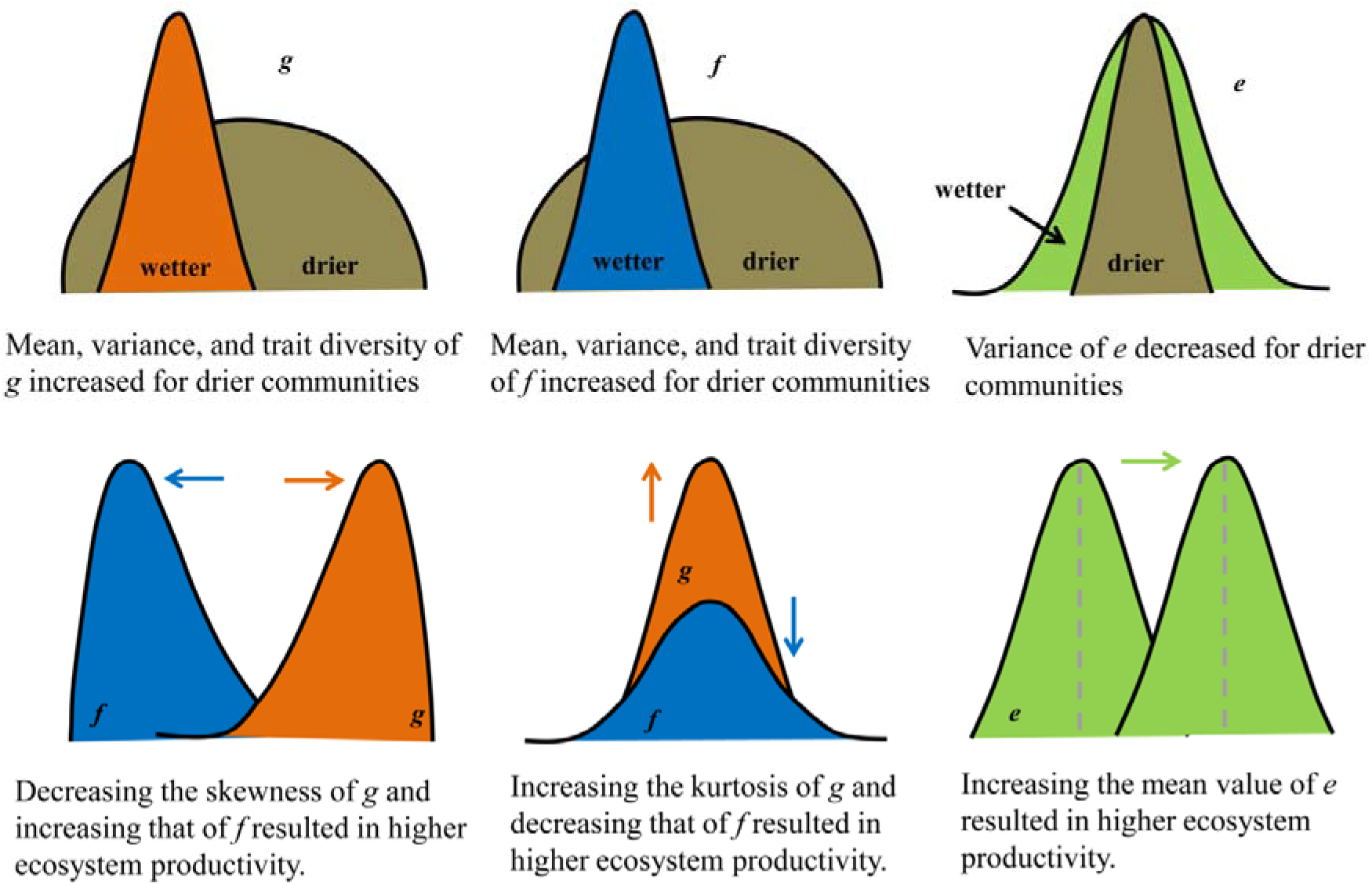
Conceptual diagrams of how stomatal trait distributions adapt drought stress and regulate ecosystem productivity. *g*, maximum stomatal conductance, shown by orange; *f*, stomatal area fraction, shown by blue; *e*, stomatal space-use efficiency, shown by green.

Overall stomatal traits explained up to 66% of the total variation observed in ecosystem productivity, which was greater than that explained by the distributions of stomatal traits generated by the two null models (Fig. S2).

## Discussion

### Maximum stomatal conductance (*g*) increases with climatic aridity at continental scale

The linkage of *g* with low water availability has remained controversial. Indeed, plants may adapt to dry conditions with a low *g* that may enable sustained low rates of gas exchange under extended periods of lower water supply, with increased CO_2_ gain relative to water loss, i.e., higher water use efficiency (Franks et al., 2015). However, some studies have proposed a higher *g* and stomatal conductance can confer an advantage for plants in arid climates, enabling greater rates of photosynthesis in the shorter “pulses” when water is available (Grubb, 1998; Scoffoni et al., 2011; Wang et al., 2017), and thus “avoiding” drought with opportunistic rapid growth during short periods of water availability. One of the major novel findings of this study was that the community-weighted mean value of *g* was positively related to climatic aridity across the continent, and thus that pulse-driven “avoidance” is the dominant trend for adaptation of communities with low water availability. Our findings extend to continental scale the hypothesis that plants and communities adapted to arid climates would generally maintain a low stomatal conductance, but given their high maximum stomatal conductance, can sharply increase stomatal conductance during pulses of rainfall availability to maximize growth (Grubb, 1998). This hypothesis is also consistent with reports that species with higher *g* tend to show greater sensitivity to changes in the external environment (Haworth et al., 2018; Siddiq et al., 2017).

### Greater functional niche differentiation of *g* under higher climatic aridity

Environmental stress can restrict the variance of trait values, leading to convergence in the distribution of trait values among coexisting species (Kraft et al., 2008). Yet, for communities across a continental scale aridity gradient, the community-weighted variance of *g* increased with climatic aridity. Notably, this difference for *g* would be expected due to the ability of plants to close stomata; species with a high *g* are not obliged to maintain high stomatal conductance during stressful periods, as species with large stomatal pores areas can sharply reduce stomatal conductance and thus transpiration rates. The distance between observed kurtosis and minimum kurtosis (D) for *g* was lower than that generated by two null models (Table S1), consistent with a general assembly rule that trait diversity of *g* is maximized within forest plant communities, as previously demonstrated for global drylands in analyses of specific leaf area and maximum height (Gross et al., 2017). Further, the D values of *g* was lower for drier communities, suggesting that this assembly rule applies more strongly with increasing aridity. Similarly to root stratification (Oram et al., 2018), diversity in *g* and associated stomatal regulation strategies might improve species-specific soil moisture status (West et al., 2012) and increase species partitioning water resources in space and/or time, thus increasing overall water utilization (Naeem et al., 1994). We observed a negative relationship between community-weighted variance and kurtosis of *g* (Fig. S1); communities characterized by low variance and low kurtosis values were only observed in the wetter regions, indicating that community assembly process of *g* was more strictly constrained under lower water availability. The strong patterns linking the stomatal traits of communities with climate at continental scale highlights the importance of these traits across the background of other structural and physiological adaptations to aridity, including specialized xylem anatomy, plant allometry, rooting strategy, dormancy and the ability to recover after dieback (Grossiord, 2020).

### Limited variability of stomatal space-use efficiency (*e*) under water scarcity

Stomatal space-use efficiency (*e*) was first defined in this study, and, by contrast with *g* and *f*, community-weighted mean values of *e* were not statistically constrained by climatic aridity, supporting theory that this efficiency should be generally maximized (FranksandBeerling, 2009). For *g* and *f*, the overall negative correlation community-weighted mean trait values with aridity was consistent with the expected trends based on adaptation (GarnierandNavas, 2012; Garnier et al., 2004; Grime, 1998). Likewise, given that community-weighted mean values of *e* were highly conservative, the narrow community-weighted variance of *e* would reflect adaptation in which co-occurring species tend to converge in *e* to a narrow range of optimal values. Our results supported the hypothesis that the variability of *e* was especially strongly constrained under arid climates, consistent with the expectation of greater cost-effectiveness of investment in stomata under lower water availability than under high water availability, where selection would likely be weaker.

### Coordinated adaptation of *g* and *f* across a climatic gradient

For both *g* and *f*, the community-weighted mean and variance increased with the climatic aridity, whereas D decreased, and the trait diversity was maximized. Thus, the distributions of *g* and *f* were synchronous in adapting to the environment. Given that *g* is determined as the product of *f* and *e*, and that variation in *g* was primarily caused by *f* rather than *e*, it is clear that the shifts in stomatal area fraction are more typical for the adaptation and assembly of *g* than shifts in *e*, which remains constrained. As *e* is inversely proportional to stomatal size (see Supplementary Note 1 for detailed information), its constraint is consistent with previous studies reporting that stomatal size is less variable than stomatal density or *f* (Beaulieu et al., 2008; Jordan et al., 2015; XiongandFlexas, 2020).

### Contrasting roles of *g* and *f* in optimizing ecosystem productivity

Selection for higher *g* (the benefit) involves a trade-off to minimize *f* (the cost) (de Boer et al., 2016), and such cost-benefit relationship is also involved in how stomatal traits regulate ecosystem productivity. Decreasing the skewness of *g* and increasing the skewness of *f* meant that species with high *g* and/or low *f* values were more dominant within communities; thus, the optimization of stomata on ecosystem productivity was economical through decreasing the skewness of *g* and increasing the skewness of *f*. A previous study also argued that high kurtosis in leaf traits indicated strong trait optimization (Umaña et al., 2021). Here, the high kurtosis of *g* meant that co-occurring species of *g* were convergent toward an optimal value. Nevertheless, the high kurtosis and lower skewness of *g* coupled with lower kurtosis and higher skewness of *f* would result in improved *e*, i.e., the benefit-cost ratio (de Boer et al., 2016), which was positively correlated with ecosystem productivity. Therefore, contrasting regulations of *g* and *f* on ecosystem productivity were associated with stomatal cost-benefit relationship.

## Materials and Methods

### Study sites and climate data

Nine study sites were selected along the 3700-km north–south transect of China (NSTEC), which were designated as Huzhong, Liangshui, Changbai, Dongling, Taiyue, Shennongjia, Jiulian, Dinghu, and Jianfengling. The nine study sites extend from 18.7 °N to 51.8 °N latitude, and represent examples of most of the forest types in the northern hemisphere, including cold-temperate coniferous forest, temperate deciduous forest, subtropical evergreen forest, and tropical rain forest (He et al., 2019). Along the NSTEC transect, the mean annual temperature (MAT) ranges from –3.67 to 23.2 °C, and mean annual precipitation (MAP) from 472 to 2266 mm (He et al., 2020). Soil types range from cold-temperate brown soils with high organic matter content to tropical red soils with low organic matter content.

### Sample collection and analysis

The field survey was conducted in July–August 2013, the peak period of growth for all species. Sampling plots were located within well-protected national nature reserves with relatively continuous vegetation, which is representative of the given forests. Three or four experimental plots (30 m × 40 m) located least 100 m apart were established in each site. Geographical information (latitude, longitude, and altitude), plant species composition, and community structure were recorded for each plot. The number, height, diameter at breast height (DBH) of trees, basal stem diameter of shrubs, and aboveground live-biomass of all herbs were measured (He et al., 2018).

Leaves were collected from trees, shrubs, and herbs within the plots. For each species, more than 20 mature leaves were collected from the top of the canopy of four healthy individuals and mixed as a composite sample. The leaves were collected from trees using long-handle shears or handpicked by climbing the trees. About half of the leaves were placed in sealed plastic bags, immediately stored in a box with ice, and others were used to measure leaf morphological traits (Li et al., 2018).

After sampling, leaf size was measured using a scanner (Cano Scan LIDE 100, Japan) and Photoshop CS software (Adobe, United States). These leaves were subsequently dried to constant mass in an oven before measuring leaf dry mass, and specific leaf area as the ratio of leaf area to leaf dry mass. Eight to ten leaves from the pooled sample were cut into small pieces (1.0 × 0.5 cm) along the main vein and were fixed in 75% alcohol: formalin: glacial acetic acid: glycerin (90:5:5:5).

Stomatal traits were imaged using a scanning electron microscope (S-3400N, Hitachi, Japan), using the same leaf samples as previously studied for stomatal density, size and stomatal area fraction (Liu et al., 2018). Three small pieces were selected from the pooled sample, and each replicate was photographed twice on the lower surface at different positions. Given our use of scanning electron microscopy and investigation of a large number of species across communities, the labor and expense did not allow measurements of the upper epidermis, and we focused on the lower epidermis (Liu et al., 2019). The herbaceous species in closed forests typically have more stomata on their adaxial surfaces, whereas trees and shrubs tend to have few or no stomata on the adaxial surface (Muir, 2015; Muir, 2018). Thus, sampling only the lower epidermis results in some uncertainty, but the community level findings are expected to be robust.

The number of stomata in each photograph was recorded, and stomatal density (SD) was calculated as the number of stomata per unit area (Liu et al., 2018). In each photograph, five typical stomata were selected to measure stomatal length (SL), stomatal pore length (PL), and stomatal width (SW) by using MIPS (Optical Instrument Co., Ltd., Chongqing, China). We used the above stomatal traits to calculate *f* and *g* (FranksandFarquhar, 2001).

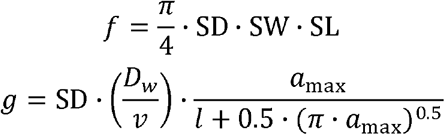

where *D*_w_ is the diffusivity of water in air, *v* is the molar volume of water vapor, *a*_max_ is the maximum pore area (estimated as the area of the ellipse with major axis PL and minor axis 0.5PL), and *l* is the depth of the stomatal pore, which was approximated as guard cell width. We then calculated *e* as the ratio of *g* to *f*. Notably, *e* depends inversely on stomatal size, because smaller stomata, having shorter depths, are more efficient for transport for a given pore area (FranksandFarquhar, 2006); the mathematical relationships of *e* to stomatal size is presented in Supplementary Note 1.

### Stomatal trait moments of plant communities

To scale up traits to the community scale, and given that stomatal traits were normalized by leaf area, we used the total leaf area of each species in the plot to weight species trait values, and then calculated the distributions of stomatal traits. The total leaf biomass of each individual tree and shrub was calculated using species-specific allometric regressions based on measured values of height, diameter at breast height (DBH) or basal stem diameter, and then the leaf biomass of each species within plots was calculated. Species-specific allometric regressions were obtained from the Chinese Ecosystem Research Network (Wang et al., 2015). The leaf biomass of herbs was measured using the harvest method. The total leaf area of each species was calculated as the product of total leaf biomass and specific leaf area. Community-weighted mean, variance, skewness, and kurtosis were calculated as follows (Gross et al., 2017; Wieczynski et al., 2019):

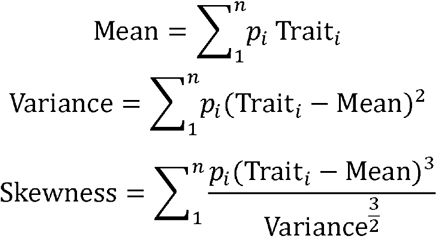

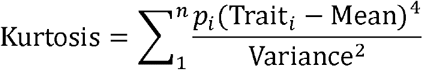

where *n* is the species richness, *p*_*i*_ is the proportion of leaf area of *i*^th^ plant species in a specific community, and Trait_*i*_ represents stomatal traits (*g, f*, or *e*) of the *i*^th^ plant species.

The community trait variance, skewness, and kurtosis provide information beyond the community weighted mean, which can over-emphasize the role of dominant species (Enquist et al., 2015). Specifically, the community variance in a given traits represents the functional divergence, skewness the extent of asymmetric distribution of traits, and kurtosis the functional evenness, with a high kurtosis indicating strong trait optimization (Umaña et al., 2021). Skewness and kurtosis are mathematically related, according to skewness-kurtosis relationships (SKR):

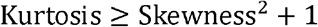

Thus, for a given skewness, there is a minimum kurtosis. Here, we calculated the distance between the observed kurtosis and minimum kurtosis (D):

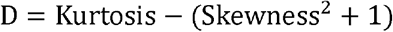

D signifies the extent to which functional diversity is maximized, with a D = 0 representing the strongest possible maximization of functional diversity (Gross et al., 2017).

### Climate data and ecosystem productivity

Mean annual temperature and precipitation (MAT and MAP, respectively) were derived from the Resource and Environment Data Cloud Platform (http://www.resdc.cn/). Then, the de Martonne aridity index (de Martonne, 1926) was calculated the ratio of MAP and MAT+10. To facilitate the interpretation of results, we calculated the climatic aridity index for each site as:

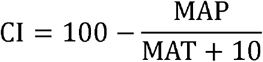

so all the CI values were positive, and higher values of this aridity level indicate drier conditions.

In these forests, gross primary productivity and net primary productivity were strongly correlated with each other across sites (Li et al., 2020); here, we focused on gross primary productivity (GPP). The average GPP data from 2000 to 2015 (Li et al., 2020) were obtained from the Numerical Terradynamic Simulation Group (http://www.ntsg.umt.edu/project/modis/mod17.php). This dataset was derived from a widely used Moderate Resolution Imaging Spectroradiometer product, and was calculated using the C5 MOD17 algorithm with data validation from flux towers (Li et al., 2020; ZhaoandRunning, 2010; Zhao et al., 2005).

### Data analysis

We calculated statistical moments for stomatal traits, including mean, variance, skewness, and kurtosis, for each of the 32 plant community plots. We tested whether to consider plots independently, rather than as nested within sites, for calculating community scale moments by comparing fixed effects models (*lm* function in R) and mixed effects models (*lmer* function from R package *lme4*). The fixed model considered plots as independent, and the mixed effects models, considered plots as a random factor nested within each site. Akaike information criterion (AIC) represented the support of the model by data, with the model having a lower AIC value more likely to underlie the data (BurnhamandAnderson, 2004). The AIC values of fixed and mixed effects models were compared, with differences greater than 2 considered decisive in selecting one model over another, representing a >100 times higher likelihood that the data were generated by that model. For 12 of the 13 relationships of traits with climate or ecosystem productivity tested in this study, the fixed effects model was selected (Table S1-S5). Thus, in our analyses, we considered each plot as a sample plant community.

Spearman rank correlation was used to test relationships between stomatal trait moments and climate variables. Ordinary least square regression was used to quantify relationships between statistical moments of stomatal traits, including the relationship between skewness^2^ and kurtosis, and relationship between variance and kurtosis. To explore whether climatic aridity mediated the relationships between variance and kurtosis, plant communities were classified into wet and dry communities (threshold CI=40), and scatter diagrams of variance and kurtosis were plotted.

Focusing on the distance to the minimal kurtosis (D) enables resolution of variation across communities in trait evenness (Gross et al., 2021), and a test of the hypothesis that functional niche differentiation of *g* would be greater in drier communities, by determining the correlation between the distance to the minimal kurtosis (D) and climatic aridity. To clarify whether trait assembly of stomata would maximize stomatal trait diversity, we tested whether observed skewness-kurtosis relationships (SKRs) differed from random expectations, which can reveal the signature of niche differentiation in shaping ecological communities (Gross et al., 2021). We constructed two null models, and predictions from each null model were derived from 2000 randomizations. In the first null model, we randomized the stomatal traits across all species, using the function “richness” in the R package PICANTE (Kembel et al., 2010). In the second null model, we shuffled stomatal traits across species occurring in each community, using the function “independentswap” in the R package PICANTE. These two null models have been the most common for analyzing community assembly, with the second null model more specific in its implication. The first null model allows tests for maximizing trait diversity locally, relative to a scenario of random selection of species from the regional pool. The second null model allows tests for maximizing trait diversity locally relative to a scenario of random selection of species from local pools. Stomatal trait moments were then calculated for each of the 2000 randomizations, for each of the null models used. Then, we assessed whether the observed SKR significantly differed from SKR_random_.Monte Carlo analysis was used to test whether the observed SKRs differed from random expectations. We compared the observed slope β and intercept α (β_obs_ and α_obs_, respectively) of the SKR with those generated by null models (β_random_ and α_random_, respectively). Three Pseudo P values were calculated: P (β|α), the frequency of β_obs_ > β_random_ within subset α_obs_ < α_random_; P (α|β), the frequency of α_obs_ > α_random_ within subset β_obs_ < β_random_; and P (β∩α), the frequency of α_obs_ < α_random_ within subset β_obs_ < β_random_. Further, we compared the observed distance to the minimal kurtosis (D_obs_) with that generated by null models (D_random_). P(D) is the frequency of D_obs_ < D_random_.

A multiple regression model was used to assess the potential influence of stomatal trait moments on ecosystem productivity, and quadratic terms of stomatal trait moments were also considered as potential drivers of non-linear effects of these variables on ecosystem productivity. All variables, including ecosystem productivity and stomatal trait moments, were standardized (Z-scores) before analysis. We first used the “stepAIC” function (MASS package in R) to exclude less important predictors, then the “dredge” function (MuMIn package in R) was used to select the best models. Finally, the relative effect of each stomatal trait moment on ecosystem productivity was calculated as its absolute parameter compared with the sum of all the absolute parameters in the model.

Standardized effect sizes (SES) were used to assess the non-random influence of stomatal traits on ecosystem productivity. SES was calculated as (Bruelheide et al., 2018)

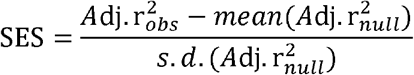

where 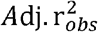 is the observed influence of stomatal traits on ecosystem productivity, 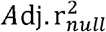 is the influence of stomatal traits on ecosystem productivity of random communities generated from a null model, *mean* represents the average value, and *s*.*d*. is the standard deviation.

Data analyses and visualization were performed using R (http://www.R-project.org/).

Statistical significance was set at the 0.05 level.

## Author contribution

N.H. planned and designed the research; C.L. and Y.L. conducted fieldwork and collected data; C.L., L.S. and Y.L. analyzed data and wrote the manuscript; L.S. and C.L. revised the manuscript.

## Acknowledgements

This work was supported by National Natural Science Foundation of China [31988102, 31770655, 32001186], and the fellowship of China Postdoctoral Science Foundation (2020M680663). There are no conflicts of interest to declare.

## Data accessibility

The data that support the findings of this study are available from the corresponding author upon reasonable request.

