## Supplementary material for "Contrasting adaptation and optimization of stomatal traits across communities at continental-scale": SI

**Table S1. Comparison of fixed effects and mixed effects models for testing relationships between community-weighted mean (CWM) of stomatal traits and climatic aridity (CI)**

| Model | Formula | $R^2_m$ | $R^2_c$ | AIC |
|---|---|---|---|---|
| Fixed effects model★★ | lm(log(CWM_*g*, base=10)~CI) | 0.58 | / | -28.60 |
| Mixed effects model | lmer(log(CWM_*g*,base=10)~CI+(1\|Site)) | 0.52 | 0.82 | -22.06 |
| Fixed effects model★ | lm(log(CWM_*f*, base=10)~CI) | 0.49 | / | -33.77 |
| Mixed effects model★ | lmer(log(CWM_*f*,base=10)~CI+(1\|Site)) | 0.45 | 0.84 | -32.01 |
| Fixed effects model★★ | lm(log(CWM_*e*, base=10)~CI) | 0.02 | / | -59.80 |
| Mixed effects model | lmer(log(CWM_*e*,base=10)~CI+(1\|Site)) | 0.01 | 0.67 | -55.23 |

$R^2_m$, marginal $R^2$; $R^2_c$, conditional $R^2$. The marginal R-squared describes the variation explained by the fixed factor only, whereas the conditional R-squared describes the variation explained by the fixed and random factors together.

AIC, Akaike information criterion.

★★ indicates the model is selected with AIC lower by 2; ★ indicates that the AIC values for the different models are within 2 of each other.

*g*, maximum stomatal conductance; *f*, stomata area fraction; *e*, stomatal space-use efficiency.

**Table S2 Comparison of fixed effects and mixed effects models for testing relationships between community-weighted variance (CWV) of stomatal traits and climatic aridity (CI)**

| Model | Formula | $R^2_m$ | $R^2_c$ | AIC |
|---|---|---|---|---|
| Fixed effects model★ | lm(log(CWV_*g*, base=10)~CI) | 0.66 | / | 8.17 |
| Mixed effects model★ | lmer(log(CWV_*g*,base=10)~CI+(1\|Site)) | 0.60 | 0.88 | 8.12 |
| Fixed effects model | lm(log(CWV_*f*, base=10)~CI) | 0.40 | / | 22.94 |
| Mixed effects model★★ | lmer(log(CWV_*f*,base=10)~CI+(1\|Site)) | 0.37 | 0.82 | 19.57 |
| Fixed effects model★★ | lm(log(CWV_*e*, base=10)~CI) | 0.37 | / | 15.36 |
| Mixed effects model | lmer(log(CWV_*e*,base=10)~CI+(1\|Site)) | 0.35 | 0.46 | 30.09 |

$R^2_m$, marginal $R^2$; $R^2_c$, conditional $R^2$. The marginal R-squared describes the variation explained by the fixed factor only, whereas the conditional R-squared describes the variation explained by the fixed and random factors together.

AIC, Akaike information criterion.

★★ indicates the model is selected with AIC lower by 2; ★ indicates that the AIC values for the different models are within 2 of each other.

*g*, maximum stomatal conductance; *f*, stomata area fraction; *e*, stomatal space-use efficiency.

**Table S3 Comparison of fixed effects and mixed effects models for testing relationships between community-weighted skewness (CWS) and community-weighted kurtosis (CWK)**

| Model | Formula | $R^2_m$ | $R^2_c$ | AIC |
|---|---|---|---|---|
| Fixed effects model★★ | lm(CWK_*g*~CWS_*g*$^2$) | 0.75 | / | 125.1 |
| Mixed effects model | lmer( CWK_*g*~CWS_*g*$^2$+(1\|Site)) | 0.72 | 0.80 | 127.4 |
| Fixed effects model★★ | lm( CWK_*f*~CWS_*f*$^2$) | 0.77 | / | 95.6 |
| Mixed effects model | lmer( CWK_*f*~CWS_*f*$^2$+(1\|Site)) | 0.72 | 0.80 | 98.0 |
| Fixed effects model★★ | lm( CWK_*e*~CWS_*e*$^2$) | 0.86 | / | 138.4 |
| Mixed effects model | lmer( CWK_*e*~CWS_*e*$^2$+(1\|Site)) | 0.86 | 0.90 | 141.0 |

$R^2_m$, marginal $R^2$; $R^2_c$, conditional $R^2$. The marginal R-squared describes the variation explained by the fixed factor only, whereas the conditional R-squared describes the variation explained by the fixed and random factors together.

AIC, Akaike information criterion.

★★ indicates the model is selected with AIC lower by 2; ★ indicates that the AIC values for the different models are within 2 of each other.

*g*, maximum stomatal conductance; *f*, stomata area fraction; *e*, stomatal space-use efficiency.

CWS, community-weighted skewness; CWK, community-weighted kurtosis

**Table S4 Comparison of fixed effects and mixed effects models for testing relationships between the distance between observed kurtosis and minimum kurtosis (Distance) and climatic aridity index (CI)**

<table>
<tr><th>Model</th><th>Formula</th><th>$R^2_m$</th><th>$R^2_c$</th><th>AIC</th></tr>
<tr><td>Fixed effects model★★</td><td>lm(log(Distance_g, base=10)~CI)</td><td>0.52</td><td>/</td><td>30.63</td></tr>
<tr><td>Mixed effects model</td><td>lmer(log(Distance_g,base=10)~CI+(1|Site))</td><td>0.48</td><td>0.71</td><td>40.45</td></tr>
<tr><td>Fixed effects model★★</td><td>lm(log(Distance _f, base=10)~CI)</td><td>0.44</td><td>/</td><td>19.28</td></tr>
<tr><td>Mixed effects model</td><td>lmer(log(Distance_f,base=10)~CI+(1|Site))</td><td>0.42</td><td>0.54</td><td>33.14</td></tr>
<tr><td>Fixed effects model★★</td><td>lm(log(Distance_e, base=10)~CI)</td><td>0.05</td><td>/</td><td>17.68</td></tr>
<tr><td>Mixed effects model</td><td>lmer(log(Distance_e,base=10)~CI+(1|Site))</td><td>0.04</td><td>0.39</td><td>29.53</td></tr>
</table>

$R^2_m$, marginal $R^2$; $R^2_c$, conditional $R^2$. The marginal R-squared describes the variation explained by the fixed factor only, whereas the conditional R-squared describes the variation explained by the fixed and random factors together.

AIC, Akaike information criterion.

★★ indicates the model is selected with AIC lower by 2; ★ indicates that the AIC values for the different models are within 2 of each other.

*g*, maximum stomatal conductance; *f*, stomata area fraction; *e*, stomatal space-use efficiency.

**Table S5 Comparison of fixed effects and mixed effects models for testing relationships between stomatal trait moment and ecosystem productivity (GPP).**

| Model | Formula | AIC |
|---|---|---|
| **Fixed effects model★★** | lm(GPP~CWM_g+CWV_g+CWS_g+CWK_g+CWM_f+CWV_f+CWS_f +CWK_f+CWM_e+CWV_e+CWS_e+CWK_e+I(CWM_g^2)+I(CWV_g^2) +I(CWS_g^2)+I(CWK_g^2)+I(CWM_f^2)+I(CWV_f^2)+I(CWS_f^2) +I(CWK_f^2)+I(CWM_e^2)+I(CWV_e^2)+I(CWS_e^2)+I(CWK_e^2)) | 65.73 |
| **Mixed effects model** | lmer(GPP~CWM_g+CWV_g+CWS_g+CWK_g+CWM_f+CWV_f +CWS_f+CWK_f+CWM_e+CWV_e+CWS_e+CWK_e +I(CWM_g^2)+I(CWV_g^2)+I(CWS_g^2)+I(CWK_g^2) +I(CWM_f^2)+I(CWV_f^2)+I(CWS_f^2)+ I(CWK_f^2)+I(CWM_e^2)+I(CWV_e^2) +I(CWS_e^2)+I(CWK_e^2)+(1\|Site)) | 116.72 |

$R^2_m$, marginal $R^2$; $R^2_c$, conditional $R^2$. The marginal R-squared describes the variation explained by the fixed factor only, whereas the conditional R-squared describes the variation explained by the fixed and random factors together.

AIC, Akaike information criterion.

★★ indicates the model is selected with AIC lower by 2; ★ indicates that the AIC values for the different models are within 2 of each other.

*g*, maximum stomatal conductance; *f*, stomata area fraction; *e*, stomatal space-use efficiency.

CWM, community-weighted mean; CWV, community-weighted variance; CWS, community-weighted skewness; CWK, community-weighted kurtosis.

**Table S6 Results from the null model for stomatal traits**

| Direct assessments | | | | | | | | | Distance to the lower boundary | | |
|---|---|---|---|---|---|---|---|---|---|---|---|
| **Traits** | **Observed** | **$R^2$** | **Null model** | **P(β)** | **P(α)** | **P(β\|α)** | **P(α\|β)** | **P(β∩α)** | **Traits** | **Null model** | **P(D)** |
| ***g*** | β = 1.87 | 0.74 | Richness | 0.936 | 0.017 | 0.029 | 0.001 | < 0.001 | ***g*** | Richness | < 0.001 |
| | α = 1.39 | | Swap | 0.958 | 0.048 | 0.463 | 0.023 | 0.022 | | Swap | 0.361 |
| **log (*g*)** | β = 1.54 | 0.85 | Richness | 0.681 | 0.022 | < 0.001 | < 0.001 | < 0.001 | **log (*g*)** | Richness | < 0.001 |
| | α = 2.26 | | Swap | 0.543 | 0.113 | 0.013 | 0.003 | 0.001 | | Swap | 0.013 |
| ***f*** | β = 1.54 | 0.76 | Richness | 0.720 | 0.065 | 0.031 | 0.003 | 0.002 | ***f*** | Richness | < 0.001 |
| | α = 1.90 | | Swap | 0.738 | 0.351 | 0.408 | 0.194 | 0.143 | | Swap | 0.061 |
| **log (*f*)** | β = 1.36 | 0.41 | Richness | 0.577 | 0.277 | 0.260 | 0.125 | 0.072 | **log (*f*)** | Richness | < 0.001 |
| | α = 2.83 | | Swap | 0.565 | 0.420 | 0.277 | 0.206 | 0.116 | | Swap | < 0.001 |
| ***e*** | β = 1.38 | 0.86 | Richness | 0.427 | 0.782 | 0.328 | 0.601 | 0.256 | ***e*** | Richness | 0.602 |
| | α = 3.46 | | Swap | 0.551 | 0.519 | 0.272 | 0.256 | 0.141 | | Swap | 0.366 |
| **log (*e*)** | β = 1.10 | 0.90 | Richness | 0.296 | 0.718 | 0.193 | 0.468 | 0.139 | **log (*e*)** | Richness | 0.446 |
| | α = 3.14 | | Swap | 0.270 | 0.550 | 0.114 | 0.232 | 0.063 | | Swap | 0.217 |

*g*, maximum stomatal conductance; *f*, stomatal area fraction; *e*, stomatal space-use efficiency.
Direct assessments of probabilities derived from the null models for the skewness-kurtosis relationships (SKRs). The parameters intercept (β) and slope (α) were tested separately, conditionally and jointly under the two randomization procedures. P(β), probability of finding a lower random slope β during the randomization procedure than the observed slope β; P(α), probability of finding a lower random y-intercept α than the observed α; P(β|α), conditional probability of finding a lower random slope than the observed slope; P(α|β), conditional probability of finding a lower intercept in the randomizations than observed; P(β∩α), probability of finding in the randomizations both a lower intercept and slope than observed.
We indicate the regression parameters for observed SKRs. Pseudo P values below 0.05 indicate that the observed parameters are lower than the parameters expected by chance.
We compared the distance (D) of observed data and the predictions of the two null models for three stomatal traits. P values below 0.05 indicate that observed values are closer to the lower boundary of the SKR than expected by chance

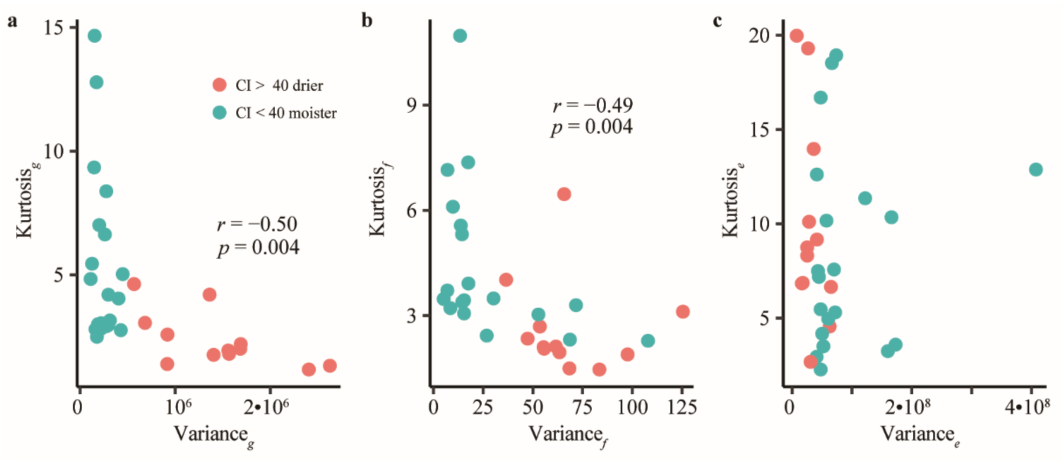


**Fig. S1 Relationships between community-weighted variance and community-weighted kurtosis of stomatal traits.**

*g*, maximum stomatal conductance; *f*, stomatal area fraction; *e*, stomatal space-use efficiency. Variance, community-weighted variance; Kurtosis, community-weighted kurtosis.

Note that here we split the communities into two groups: drier communities (aridity index> 40) and moister communities (aridity index < 40).

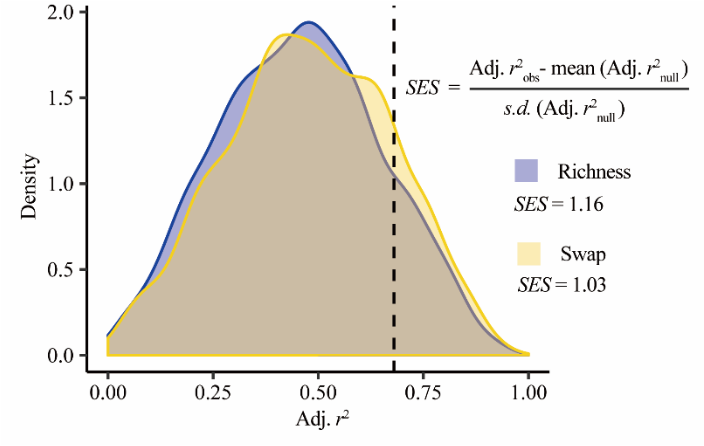


**Fig. S2 Observed effect of stomatal traits on ecosystem productivity (Adj.$r^2$) and null expectations.**

SES, standardized effect size.

Dashed line represented observed Adj.$r^2$.

In this study, we ran two null models, named “Richness” and “Swap”, see **Method** for details.
