## Supplementary material for "Contrasting adaptation and optimization of stomatal traits across communities at continental-scale": Note 1

The theoretical $\boldsymbol{g}$ was first derived from stomatal anatomical traits by Brown and Escombe (1900), and the "double end-correction equation" is the most widely used formula in recent years (Franks & Beerling 2009):

$$\boldsymbol{g} = \boldsymbol{d} \cdot \left(\frac{\boldsymbol{D_w}}{\boldsymbol{v}}\right) \cdot \frac{\boldsymbol{a}_{\mathbf{max}}}{\boldsymbol{l}+\mathbf{0.5}\cdot(\boldsymbol{\pi}\cdot\boldsymbol{a}_{\mathbf{max}})^{\mathbf{0.5}}} \qquad (1)$$

where $D_w$ is diffusivity of water in air, $v$ is the molar volume of gas, $a_{max}$ is the maximum pore area, and $l$ is the depth of the stomatal pore, which is approximated as a guard cell width, given that the fully inflated guard cell is approximately a disc in cross-section (Franks & Farquhar 2007).

Sack and Buckley (2016) showed Equation 1 could be rewritten as a function of stomatal density ($d$), size ($s$), and two scalars which depend on biophysics (b = $D_w/v$) and morphology (m), such that

$$\boldsymbol{g} = \mathbf{b} \cdot \mathbf{m} \cdot \boldsymbol{d} \cdot \boldsymbol{s}^{\mathbf{0.5}} \qquad (2)$$

where $\mathrm{m} = \frac{\pi\cdot c^2}{j^{0.5}\cdot(4\cdot h\cdot j+\pi\cdot c)}$, with $c$ was the ratio of stomatal pore length to stomatal length, $j$ was the ratio of stomatal width to stomatal length, and $h$ was the ratio of pore depth to stomatal width.

Stomatal area fraction ($\boldsymbol{f}$) is an important anatomical constrain of $\boldsymbol{g}$. Liu *et al.* (2018) calculated $f$ as follows:

$$\boldsymbol{f} = \boldsymbol{d} \cdot \boldsymbol{s} \qquad (3)$$

Applying Equation 3 to Equation 2 gives

$$\boldsymbol{g} = \mathbf{b} \cdot \mathbf{m} \cdot \boldsymbol{f} \cdot \boldsymbol{s}^{-\mathbf{0.5}} \qquad (4)$$

de Boer *et al.* (2016) highlighted that $\boldsymbol{f}$ was the cost associated with the operation and maintenance of the stomata, whereas $\boldsymbol{g}$ represented the ability of plants to assimilate carbon. Here, a trade-off exists that can maximize benefits while minimizing costs; therefore, we defined a new stomatal trait−stomatal space-use efficiency ($\boldsymbol{e}$) as follows:

$$\boldsymbol{e} = \frac{\boldsymbol{g}}{\boldsymbol{f}} \qquad (5)$$

Equation 5 can be rearranged as follows:

$$\boldsymbol{g} = \boldsymbol{f} \cdot \mathbf{e} \qquad (5a)$$

Applying Equation 4 to Equation 5 gave the following:

$$\boldsymbol{e} = \mathbf{b} \cdot \mathbf{m} \cdot \boldsymbol{s}^{-0.5} \qquad (6)$$

Under the same $\boldsymbol{f}$, combinations of higher stomatal density and smaller size would yield higher *g*; therefore, generally small stomatal size denotes higher $\boldsymbol{e}$ (Franks & Beerling 2009; de Boer *et al.* 2016). Equation 6 also showed that there should be a negative relationship between *s* and *e*. Changes in stomatal size among different plant functional groups have been tested, and many studies have demonstrated that the stomata of herbs are larger and sparser, whereas those of trees are smaller and denser (Salisbury 1928; Wang *et al.* 2015; Liu *et al.* 2018)
